# Using Mesoscopic Tract-Tracing Data to Guide the Estimation of Fiber Orientation Distributions in the Mouse Brain from Diffusion MRI

**DOI:** 10.1101/2022.06.02.492838

**Authors:** Zifei Liang, Tanzil Mahmud Arefin, Choong H. Lee, Jiangyang Zhang

## Abstract

Diffusion MRI (dMRI) tractography is the only tool for non-invasive mapping of macroscopic structural connectivity over the entire brain. Although it has been successfully used to reconstruct large white matter tracts in the human and animal brains, the sensitivity and specificity of dMRI tractography remained limited. Especially, the fiber orientation distributions (FODs) estimated from dMRI signals, key to tractography, may deviate from histologically measured fiber orientation in crossing fibers and gray matter regions. In this study, we demonstrated that a deep learning network, trained using mesoscopic tract-tracing data from the Allen Mouse Brain Connectivity Atlas, was able to improve the estimation of FODs from mouse brain dMRI data. Tractography results based on the network generated FODs showed improved specificity while maintaining sensitivity comparable to results based on FOD estimated using a conventional spherical deconvolution method. Our result is a proof-of-concept of how mesoscale tract-tracing data can guide dMRI tractography and enhance our ability to characterize brain connectivity.

## Introduction

The introduction of diffusion MRI (dMRI) based tractography more than two decades ago generated great excitement due to its promise of tracing white matter tracts in the brain non-invasively (Basser and Jones, 2002; Jbabdi et al., 2015; Mori and van Zijl, 2002). Aided by increasingly sophisticated dMRI acquisition schemes and tractography methods (Behrens et al., 2007; Frank, 2001; Mori and van Zijl, 2002; Tournier et al., 2007; Tuch et al., 2002; Wedeen et al., 2012; Wedeen et al., 2008), dMRI tractography had gained the ability to resolve multiple axonal bundles within a voxel as well as high sensitivity to small white matter tracts than the original diffusion tensor imaging-based tractography. To this date, dMRI tractography remains the only tool for non-invasive mapping of macroscopic structural connectivity of the entire brain.

Current routine dMRI tractography starts with the estimation of the fiber orientation distribution (FOD) in each voxel from dMRI signals (Tournier et al., 2007), followed by generation of streamlines connecting neighboring voxels based on FODs using either deterministic or probabilistic algorithms (Behrens et al., 2007; Mori and van Zijl, 2002). While numerous studies in human and animal brains have demonstrated the capabilities of dMRI tractography in visualizing tract trajectories and estimating structural connectivity between brain regions under normal and pathological conditions, e.g. (Catani et al., 2002; Jbabdi et al., 2015; Lo et al., 2010; White et al., 2020), its limitations have also been recognized (Jbabdi and Johansen-Berg, 2011; Jones et al., 2013), and efforts have been made to understand the underlying causes in order to circumvent or alleviate these limitations.

Numerous studies have compared tractography results with chemical and viral tracer findings from post-mortem brain specimens, e.g. (Aydogan et al., 2018; Azadbakht et al., 2015; Calabrese et al., 2015; Chen et al., 2015; Dyrby et al., 2007; Grisot et al., 2021; Howard et al., 2019; Seehaus et al., 2013). While these studies demonstrated good agreements for major white matter tracts, they also revealed noticeable differences when tractography results entered superficial white matter and gray matter regions (Grisot et al., 2021; Howard et al., 2019; Maier-Hein et al., 2017; Reveley et al., 2015; Schilling et al.,2019; Thomas et al., 2014; Wu and Zhang, 2016). Several approaches to improve dMRI tractography have been explored, such as improving spatial resolution (Calabrese et al., 2014; Liebrand et al., 2020; Poot et al., 2013), more accurate estimation of FODs (De Luca et al., 2020; Jeurissen et al., 2014), and optimizing tractography parameters (Fillard et al., 2011; Gutierrez et al., 2020; Moldrich et al., 2010; Schurr et al., 2018).

Among these approaches, the accurate estimation of FOD is of particular importance. Due to its central role in directing tractography. the estimated FODs should closely follow the orientations of underlying axonal pathways. Several studies have compared FODs estimated from dMRI signals with axon orientation distributions measured using a variety of optical techniques including confocal microscopy (Schilling et al., 2016; Schilling et al., 2018), polarized light imaging (Mollink et al., 2017), and optical coherence tomography (Jones et al., 2020). These studies generally demonstrated good but not perfect agreements between the estimated FODs and underlying axonal pathways, echoing the well-recognized challenge of inferring tissue axonal organization from dMRI signals.

In this study, we investigated whether deep learning could improve the estimation of FODs from dMRI signals. While several widely used estimators of FODs are based on dMRI signal models of myelinated axons (Tournier et al., 2007), tissue microstructure in gray matter is more complex than white matter with additional cellular compartments, and our knowledge of how to model dMRI signals from gray matter remains limited. In comparison, the deep learning framework is data-driven and model-free. There have been several reports on using machine and deep learning to predict MRI-based tissue parameters, including FODs, from MRI signals (Gibbons et al., 2019; Li et al., 2021; Lin et al., 2019), bypassing time-consuming model fitting procedures. Our recent study also demonstrated that the deep learning framework can be used to generate virtual histology from MRI signals by training a network with co-registered MRI and histological data (Liang et al., 2022). With the data from the Allen Mouse Brain Connectivity Atlas (AMBCA) (Kuan et al., 2015; Lein et al., 2007; Oh et al., 2014), which contains a large collection of mouse brain tract-tracing data captured using serial two-photon microscopy, we can now access rich information on mouse brain structural connectivity at the mesoscopic level and use it to guide the estimation of FODs from dMRI signals.

## 2. Methods

### 2.1. Generation of a whole brain streamline dataset from AMBCA

The AMBCA used a fast-marching method to generate streamlines that delineate axonal trajectories in each tracer experiment at the mesoscopic level (**Fig. 1A-B**). The labelling efficiency of the viral tracer used in AMBCA was estimated to be 40-50% (McFarland et al., 2009). The streamlines from 2,764 independent experiments were downloaded from AMBCA (http://api.brain-map.org/examples/lines/ index.html) and converted from the original JSON format into the TCK format used by MRtrix (http://www.mrtrix.org) (Tournier et al., 2019). As all the tracer data have been normalized to the Allen Reference Atlas (ARA) space, the streamlines from all experiments were aggregated to form a large streamline dataset (**Fig. 1C**). As the tracer injection sites in AMBCA experiments were mostly located in one hemisphere, uneven streamline densities between the two hemispheres were observed (**Fig. 1C**). To resolve this issue, all streamlines were mirrored along the midline of the brain, and the mirrored streamlines were added to the original streamlines to form a symmetric whole brain streamline dataset (**Fig. 1D**), which contained 3,765,678 (∼ 4 million) streamlines. We called this set of streamlines from all experiments the aggregated AMBCA streamlines to distinguish them from streamlines from individual experiments single-experiment (SE) AMBCA streamlines. The aggregated AMBCA streamlines can be downloaded from https://osf.io/m98wn.

**Fig. 1:**
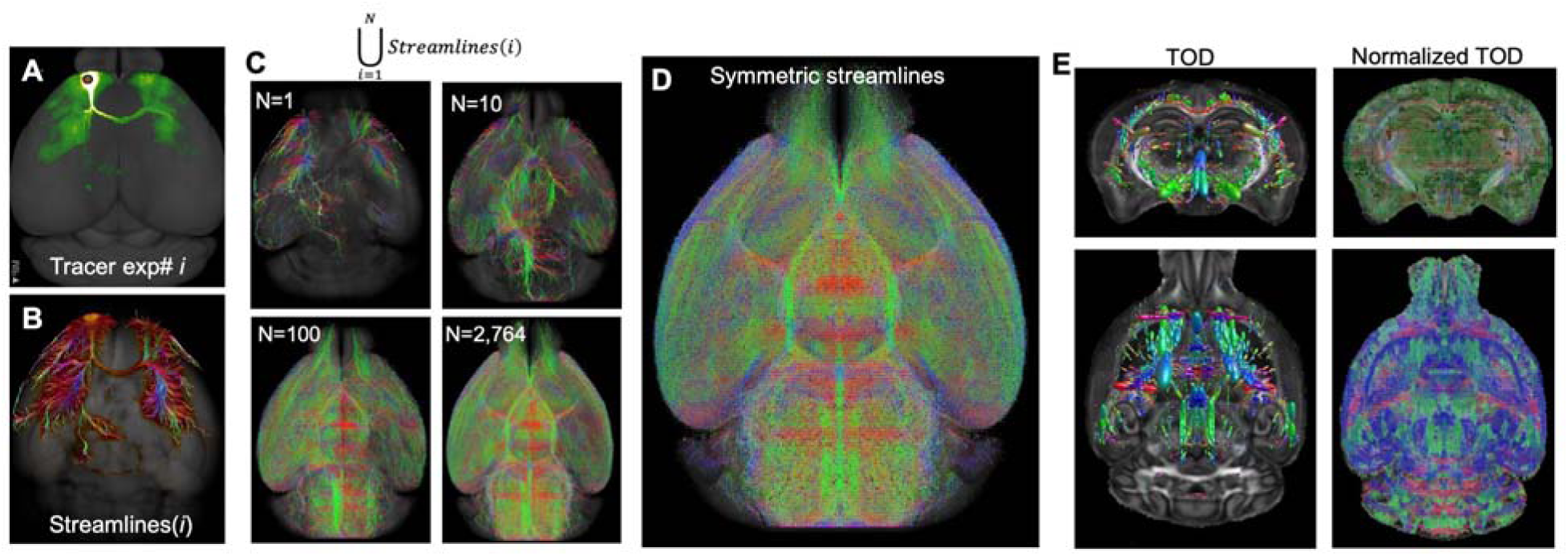
Generation of the aggregated AMBCA streamlines. **A-B:** A representative tracer experiment (ID# 100140756) and its corresponding single-experiment (SE) AMBCA streamlines generated using a fast-marching algorithm. **C:** The process of adding streamlines from all 2,764 injection experiments, with injection sites mainly located in one hemisphere. **D:** A symmetric 4-million streamline dataset was generated by mirroring the streamlines in **C** with respect to the mid-sagittal plane and added to the result in **C**. **E:** (TOD) maps generated from the aggregated AMBCA streamlines and after amplitude normalization.

### 2.2. Cell tracing data from the MouseLigtht project

To understand the differences between the mesoscopic axonal pathways from AMBCA and the path of individual axon at microscopic level, we compared the SE and aggregated AMBCA streamlines with single cell tracing data from the MouseLight project (Winnubst et al., 2019). The single-cell dataset in the MouseLight project was downloaded from its website (http://ml-neuronbrowser.janelia.org/), which contained reconstructions of 373 long-range projection neurons with axons traveling over long distances and across multiple brain regions. In this dataset, the soma, dendrites, and axons of selected neurons labelled using low-titer viral tracers were traced in serial two-photon microscopy images over the entire brain, presenting axonal pathways at the microscopic level. As the 3D two-photon image volume had been normalized to the ARA space, the downloaded axon tracing data of 373 neurons were converted from the original JSON format into the TCK format used by MRtrix (www.mrtrix.org) and gethered to form the aggregated MouseLight streamlines similar to the generation of the aggregated AMBCA streamlines but without the mirroring step.

### 2.3. Ex vivo mouse brain diffusion MRI

Mouse brain dMRI data were acquired from a separate cohort of animals. All animal experiments have been approved by the Institute Animal Care and Use Committee at New York University. Adult C57BL/6 mice (P56, n=10, 5M/5F, Charles River, Wilmington, MA, USA) were perfusion fixed with 4% paraformaldehyde (PFA) in PBS. The samples were preserved in 4% PFA for 24 hours before transferring to PBS. *Ex vivo* MRI of mouse brain specimens was performed on a horizontal 7 Tesla MR scanner (Bruker Biospin, Billerica, MA, USA) with a triple-axis gradient system. Images were acquired using a quadrature volume excitation coil (72 mm inner diameter) and a receive-only 4-channel phased array cryogenic coil. The specimens were imaged with the skull intact and placed in a syringe filled with Fomblin (perfluorinated polyether, Solvay Specialty Polymers USA, LLC, Alpharetta, GA, USA) to prevent tissue dehydration (Arefin et al., 2021). Three-dimensional diffusion MRI data were acquired using a modified 3D diffusion-weighted gradient- and spin-echo (DW-GRASE) sequence (Aggarwal et al., 2010; Wu et al., 2013) with the following parameters: echo time (TE)/repetition time (TR) = 30/400ms; two signal averages; field of view (FOV) = 12.8 mm x 10 mm x 18 mm, resolution = 0.1 mm x 0.1 mm x 0.1 mm; two non-diffusion weighted images (b0s); 60 diffusion weighted images (DWIs) with a diffusion weighting (b) of 5,000 s/mm^2^.

From the dMRI data, diffusion tensors were calculated using the log-linear fitting method implemented in MRtrix (http://www.mrtrix.org) at each pixel, and maps of mean and radial diffusivities and fractional anisotropy were generated. FOD maps were generated using constrained spherical deconvolution (CSD) implemented in MRtrix using a single response function estimated from white matter structures as described in (Arefin et al., 2021). Mappings between individual MRI data and the ARA space were computed using the Large Deformation Diffeomorphic Metric Mapping (LDDMM) (Miller et al., 2002) implemented in the DiffeoMap software (www.mristudio.org) as described in (Ceritoglu et al., 2009; Lim et al., 2013).

### 2.4. Estimation of tract-orientation-distribution (TOD) from the streamline dataset

The aggregated AMBCA streamlines were mapped from the ARA space to each subject using the mappings generated using LDDMM, which were diffeomorphic (one-to-one and differentiable) and allowed us to transform the AMBCA streamlines without changing the topology of the streamline networks. We chose this route instead of mapping dMRI data into the ARA space because the dMRI data were associated with the diffusion encoding directions used during acquisition and estimation of FODs from nonlinear mapped dMRI data requires voxel-wise adjustments of the diffusion encoding directions and amplitude, which was not a trivial task.

A 3D TOD map was then generated from the aggregated AMBCA streamlines mapped into individual subject space using the tckmap command in MRtrix with the maximum spherical harmonic degree (lmax) set to 6 (Dhollander et al., 2014). The orientation and amplitude of the TOD reflected the arrangement and number of AMBCA streamlines at each voxel after mapping. Using the initial TODs, we computed short streamlines with lengths less than 6 mm and then generated the so-called level-2 TODs based on the short streamlines as described in (Dhollander et al., 2014). As the amplitudes of TODs generated from the aggregated AMBCA streamlines depend on the distribution of injection sites and may not reflect the actual number of axons, the amplitudes of the 28 TOD coefficients were normalized at each voxel by L1 norm. We called the normalized TODs the AMBCA TODs (**Fig. 1E**).

### 2.5. Deep learning neural network to predict FODs from dMRI signals

We started with a network described in (Lin et al., 2019), which was trained to predict CSD-FODs from dMRI signals. Briefly, the network consisted of two convolutional layers (2x2x2 kernel, 1024 and 512 filters, respectively), two fully connected layers (512 and 256 nodes, respectively), and one output layer. The inputs to the network were 3x3x3 voxel patches from dMRI data as in (Lin et al., 2019), with each voxel containing 60 diffusion-weighted signals normalized by the average of the two non-diffusion-weighted signals. The outputs of the network were 28 CSD-FOD coefficients. The network was implemented in Python using Keras (www.github.com/fchollet/keras), and the training was performed on a desktop GPU (NVIDIA RTX 2080). MRI data from 10 subjects were randomly assigned to training (n=6) and testing (n=4) groups. One million 3x3x3 patches from the training mouse brain dMRI data and their corresponding CSD-FODs were used for training, within which 10% was used as the validation cohort. We used the same hyper-parameter setting as described in (Lin et al., 2019): network weights initialization by truncated normal distribution centered on 0; initial learning rate = 1e^-4^, beta1 = 0.9, beta2 = 0.999, epsilon = 1e^-08^; Momentum = 0.5; Number of epochs = 1000; Batch size = 512. We employed the commonly used Adam’s method (Duchi et al., 2011) to stochastically optimize a mean squared error loss function with some hyper-parameters assigned before training. Early stopping was also employed by monitoring whether the validation set loss stabilized within 50 epochs to prevent overfitting (Zur et al., 2009). Using the testing data as inputs to the trained network, the predicted FODs agreed well with CSD-FODs estimated from the same data using MRtrix (**Supplementary Figure 1**).

### 2.6 Deep learning neural network to predict TODs from dMRI signals

Compared to predicting CSD-FODs from dMRI data, predicting AMBCA TODs from dMRI data faced additional challenges due to the residual mismatches between MRI and AMBCA TOD data, as well as potential discrepancies at the voxel level between the AMBCA TODs, which was computed based on the number of the aggregated AMBCA streamlines and not necessarily the actual number of axons, and dMRI signals, which reflected the actual microstructural organization (e.g. axons) in the mouse brain. To address these issues, we used a modified network as shown in **Fig. 2**. Compared to the network introduced by Lin et al. (Lin et al., 2019), we kept the first two convolutional layers (2x2x2 kernel) but increased the number of filters to 1024 for both layers. We increased the number of fully connected layers from 2 to 3 with 1024, 512, and 256 nodes. We also added 40% random dropout (Demsar and Zupan, 2021; Srivastava et al., 2014; Ying, 2019) in the two convolutional layers and the first fully connected layer as a regularization during network training. In addition, we replaced the ReLU in the hidden layers with the Parametric Rectified linear unit (PReLU) that modulates the node output (y = x, when x>0; y = ax, when x<0; where *a* is a trainable parameter) to introduce nonlinear functions that can overcome the so-called dying ReLU problem (He et al., 2015). Here, we kept the 3x3x3 patch size unchanged because we assumed that the relationship between dMRI signals and axonal bundles in the same voxel should be local and independent of neighboring voxels. Different from (Lin et al., 2019), there was residual mismatches between input dMRI data and target TOD maps, which can be accommodated by the 3x3x3 patch size as suggested by our previous report (Liang et al., 2022). We increased the training data to 3 million patches to avoid over-fitting.

**Fig. 2:**
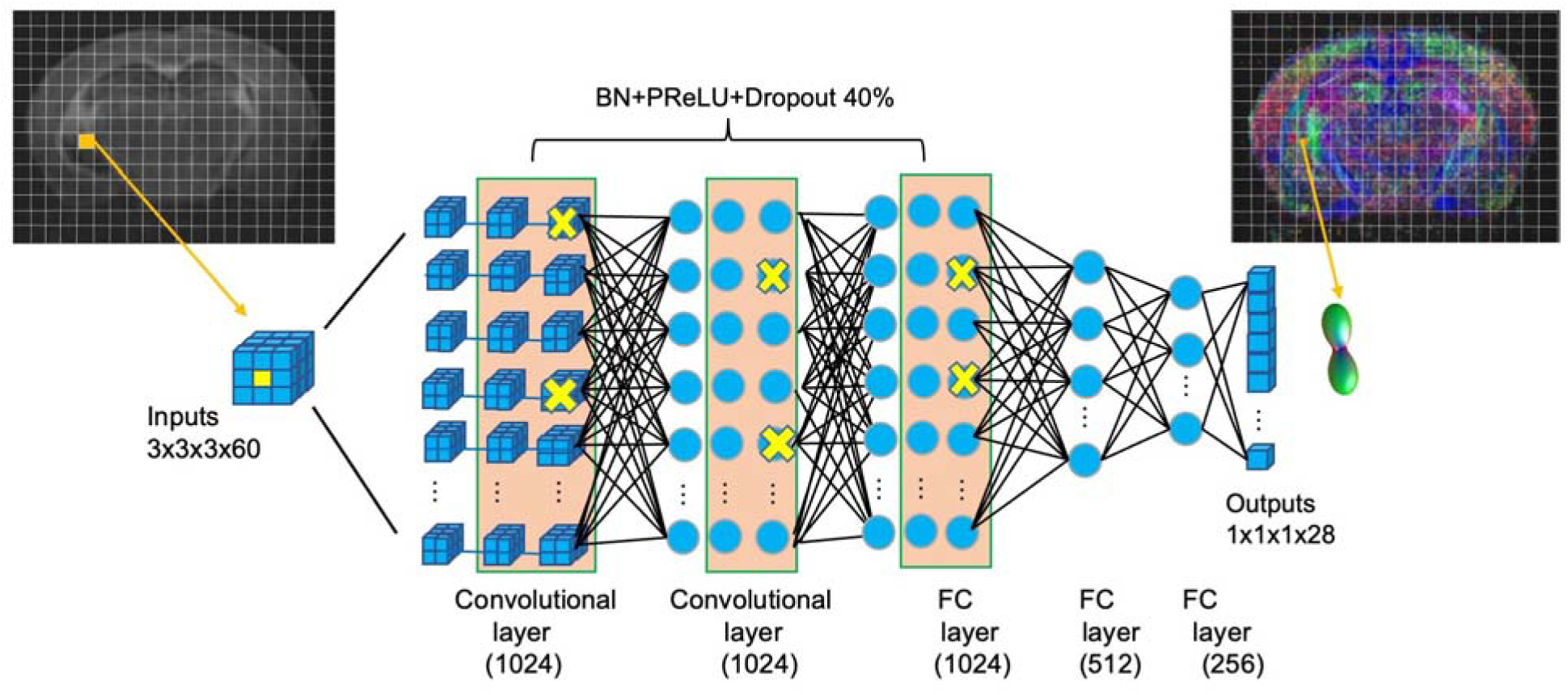
The basic architecture of TODNet. The inputs to the network were 3x3x3 voxel patches containing diffusion-weighted signals along 60 diffusion encoding directions from the voxels normalized by non-diffusion-weighted signals. The outputs were TODs with 28 spherical harmonic coefficients. The network contained 5 layers with 1024, 1024, 1024, 512, 256 output neurons, represented by the blue dots. In the first three layers, 40% neurons were dropped-off for regularization as indicated by yellow “X”.

During training, all hyper-parameters were initialized in the same way as described in section 2.5 with a mean squared error loss function. Training was carried out using an NVIDIA RTX 2080Ti and occupied 22 GB out of the total 24GB of memory with three million voxels. Since the network performed on relatively small 3x3x3 patches, it used less than 250 MB memory for voxel-wise TOD prediction. It took approximately 30 minutes to estimate a 3D DL-TOD map from a dMRI volume with a size of 256 x 200 x 300The trained models and codes are available at (https://github.com/liangzifei/MRTod_net). The output of the TODNet was called deep learning TODs (DL-TODs).

### 2.7 Tractography

Whole brain probabilistic tractography based on CSD-FODs or DL-TODs were performed using the tckgen command in MRtrix (v 3.0.3) with the FOD2 option and the optimized parameters reported in (Aydogan et al., 2018): turning angle < 45°, minimal/maximal length = 0.3/23 mm, FOD amplitude threshold = 0.1. It took approximately 2 hours to generate four million streamlines on a PC workstation (Intel Xeon E5-1650, 3.6 GHz, 64GB of RAM). The tractography streamlines were then mapped from individual subject space to the ARA space using the mappings generated in section 2.3.

From ten selected AMBCA experiments, the injection regions, which had strong intensity values, were manually defined in the tract tracing data. From the whole brain tractography streamlines mapped to the ARA space, we selected streamlines that passed through the injection regions and converted the streamline data to binary images (voxel value set to 1 if one or more streamlines passed through the voxel and 0 otherwise) for each injection region. Similarly, streamlines were selected from the aggregated AMBCA streamlines and converted to binary images, which were then dilated by 1 voxel to match the extent of axon projections in the native tracer results (**Fig. 1B**). For each subject and injection region, we used the DICE value of the binary images from tracer and tractography streamlines to quantify their spatial agreement.

### 2.8 Generation of a digital phantom

Simulated dMRI signals of crossing fibers were generated using a two-tensor model

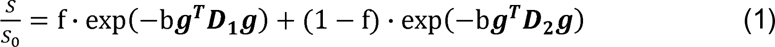

where *S* and *S0* are diffusion-weighted and non-diffusion-weighted signals, *f* the volume fraction, ***D_1_*** and ***D_2_*** the diffusion tensors, and b and ***g*** the diffusion weighting and encoding direction, respectively. In this study, we chose 12 sets of eigen values for ***D_1_*** and ***D_2_*** (**Table 1**), ranging from isotropic diffusion to highly isotropic diffusion, based on our *ex vivo* mouse brain dMRI data. A 6x6 digital phantom was created by combining different ***D_1_*** and ***D_2_*** with *f* = 0.5. We used the same diffusion weighting and diffusion encoding vectors as used in our dMRI acquisition. Simulated dMRI signals with various volume fraction (*f*) were generated using Matlab (Mathworks Inc. Natick, MA, USA) to compare estimated CSD-FODs and DL-TODs.

**Table 1:**
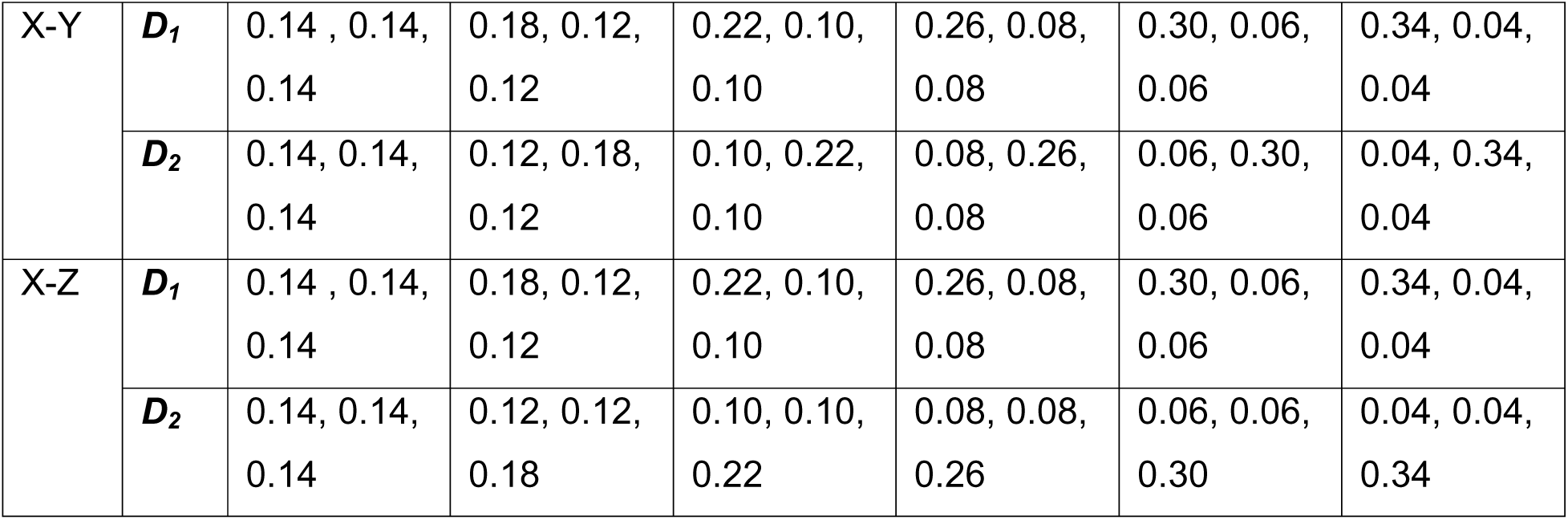
The eigen-values of ***D_1_*** and ***D_2_*** in the digital phantom for X-Y and X-Z crossing fiber configurations. The unit of the eigen-value is μm^2^/ms.

### 2.9 Statistical analysis

Two-sample paired *t*-test was used to test whether there was significant angular difference between the primary orientation of AMBCA TODs and CSD-FODs or DL-TODs from dMRI (corrections for multiple comparisons with false discovery rate = 0.1, GraphPad Prism 9.0, www.GraphPad.com) Two-sample paired *t*-test with corrections for multiple comparisons was also used to test whether there was significant difference in DICE values between binary image volumes from selected AMBCA streamlines and CSD-FOD/DL-TOD tractography results as described in section 2.7.

## 3. Results

### 3.1 The whole brain tracer streamline dataset

The aggregated AMBCA streamlines, which included data from multiple injection experiments, visualized the dense axonal pathways throughout the mouse forebrain. Major white matter tracts, such as the corpus callosum, anterior commissure, can be delineated in the dataset (**Fig. 3A**). Coherently arranged streamlines were also found in gray matter regions (e.g. cortex and hippocampus), reflecting axonal pathways in those regions, which are often challenging to reconstruct using dMRI tractography.

**Fig. 3:**
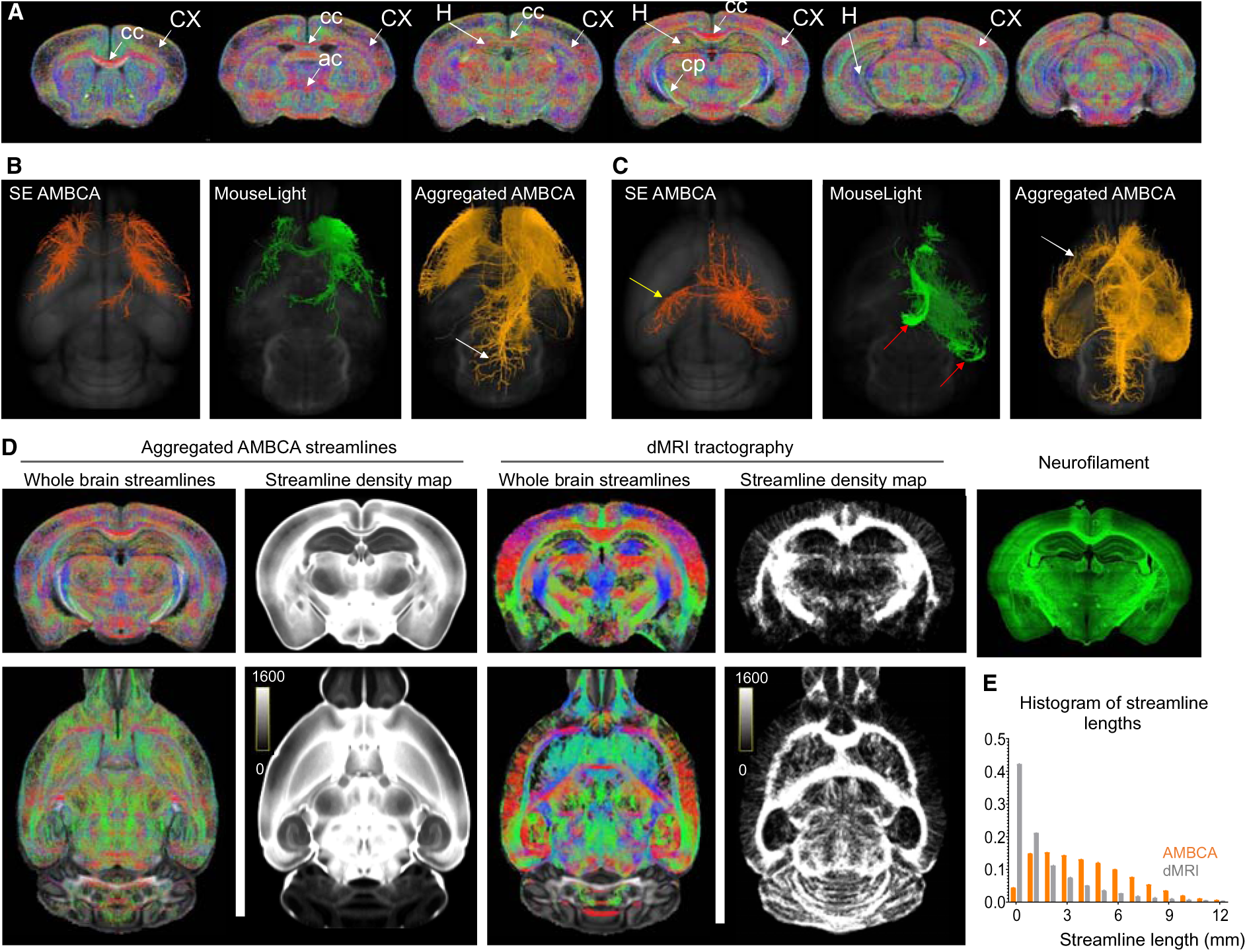
Axonal pathways in the aggregated AMBCA streamlines and comparisons with dMRI tractography. **A:** Axonal networks in the mouse brain shown in serial coronal sections of the aggregated AMBCA streamlines. Abbreviations: ac: anterior commissure; cc: corpus callosum; cp: cerebral peduncle; CX: cortex; H: hippocampus. **B-C:** A comparison of SE AMBCA streamlines originated from an injection site in the frontal cortex (B) and hippocampus (C) and streamlines in the aggregated MouseLight and AMBCA streamlines that passed the same injection site. In (C), the SE AMBCA streamlines reached the contralateral hemisphere (indicated by the yellow arrow), while the selected MouseLight streamlines showed more axonal pathways in the ipsilateral hemisphere (indicated by the red arrows). The white arrows in (B) and (C) indicate the additional streamlines found in the aggregated AMBCA streamlines. **D:** Comparisons of aggregated AMBCA and whole brain dMRI tractography streamlines acquired in this study and corresponding streamline density maps, with a matching Neurofilament-stained section from the Allen reference atlas. **E:** The histograms of streamline lengths in the AMBCA and dMRI tractography results.

We then compared SE AMBCA streamlines, which consist of anterograde streamlines originated from an injection region, with streamlines selected using the same injection region from the aggregated AMBCA and MouseLight streamlines (i.e. streamlines that passed through the injection region) (**Fig. 3B-C**). The SE AMBCA streamlines generally agreed with the selected MouseLight streamlines (e.g. **Fig. 3B**) but differences did emerged. For example, in **Fig. 3C**, the SE AMBCA streamlines showed connections to the contralateral hemisphere, whereas the selected MouseLight streamlines did not cross the midline but reached more regions within the ipsilateral hemisphere. These differences came from the fact that the MouseLight streamlines are long-range axonal projections of sparsely sampled individual neurons at the single cell level from cell tracing, whereas AMBCA streamlines represent bundles of axons at the mesoscopic level from groups of labeled neurons in the injection region.

Compared to the SE AMBCA streamlines, the selected aggregated AMBCA streamlines from the same injection region included additional streamlines that reached more regions. These additional streamlines potentially came from three sources: (1) retrograde streamlines originated from other regions but ended in the injection region; (2) streamlines that merely passed through the injection region; (3) variations in SE streamlines due to individual as well as inter-experimental variations. Compared to the selected aggregated MouseLight streamlines, the selected aggregated AMBCA streamlines from the same injection region also showed more extensive reach, and the two set of streamlines showed consistent trajectories (**Fig. 3B-C**), suggesting the aggregated AMBCA streamlines captured the major forebrain axonal pathways within the MouseLight data.

### 3.2 A comparison of the AMBCA streamline data with dMRI tractograophy

Remarkable differences were found between the aggregated AMBCA streamlines and whole brain dMRI tractography streamlines from ex vivo mouse brains. With the same number of total streamlines (4 million), the aggregated AMBCA streamline density maps showed less gray and white matter contrasts than streamline density maps from dMRI tractography (**Fig. 3D**), suggesting that tractography streamlines generated from CSD-FODs estimated here (based on dMRI data acquired here and using single white matter response function) tended to stay within white matter regions. The overall contrast pattern found in the aggregated AMBCA streamline density map resembled neurofilament stained histological sections (**Fig. 3D**), in which signal intensities reflect axonal density. In addition, the AMBCA streamlines tended to be longer than dMRI tractography streamlines (**Fig. 3E**).

### 3.3 TODNet can predict tracer streamline TOD

As expected, the DL-TOD maps generated by the TODNet based on dMRI data from the test group resembled the AMBCA TOD maps (**Fig. 4A**). In large white matter structures (e.g. the corpus callosum), the AMBCA TODs, DL-TODs, and CSD-FODs had narrow lobes along the orientations of fibers. In gray matter structures (e.g. the cortex), the CSD-FOD results showed multiple distinct lobes, whereas the AMBCA TODs and DL-TODs had no clearly defined lobes (**Fig. 4A**), making direct comparisons of DL-TODs and CSD-FODs difficult. We measured the differences in the primary orientation (the orientation of the lobe with the highest amplitude) between DL-TODs and AMBCA TODs and between CSD-FODs and AMBCA TODs at each voxel in each subject space (Fig. 4B). In white matter regions, DL-TODs were slightly better aligned to the AMBCA TODs than CSD-FODs (8.54° vs 12.94°, p<0.0001, n=4). In gray matter regions, DL-TODs showed much reduced angular difference compared to AMBCA TODs than CSD-FODs (Fig. 3C; 35.24° vs 53.16°, p<0.0001, n=4), notably in the cortex and hippocampus (**Fig. 4B**).

**Fig. 4:**
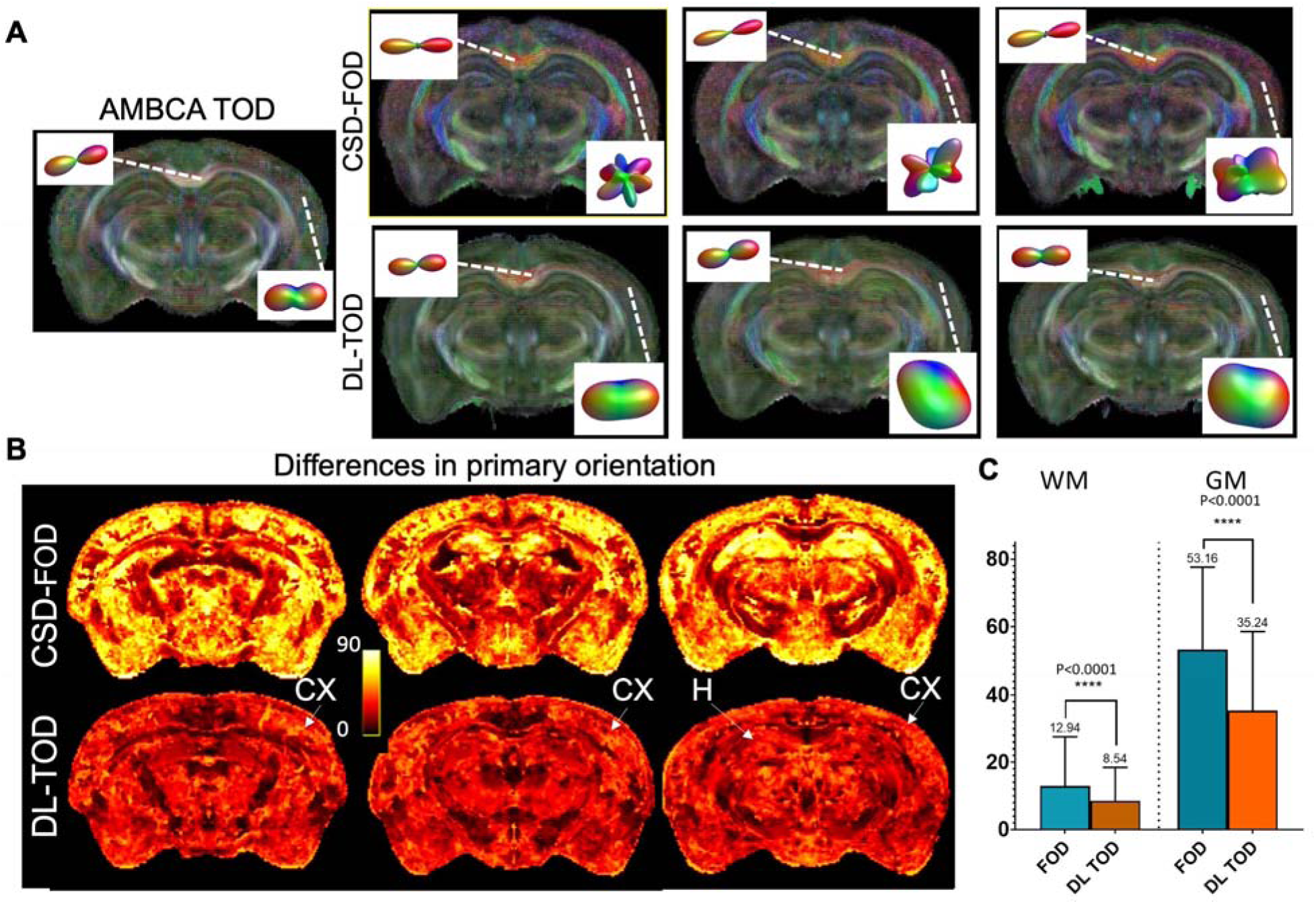
Comparisons of DL-TOD and CSD-FOD. **A:** The estimated DL-TOD and CSD-FOD maps from three mouse brains not included in training. **B:** The average differences in the primary orientation between the DL-TODs and AMBCA TODs and between the CSD-FODs and AMBCA TODs in the mouse brain data not included in training (n=4). Notice the reductions in primary orientation differences in the cortex (CX) and hippocampus (H) in the DL-TOD results compared to the CSD-FOD results. **C:** Differences in primary orientation in white matter (WM) and gray matter (GM).

We compared DL-TODs with CSD-FODs with respect to AMBCA TODs in the mouse cortex and hippocampus. In the sensory cortex, the aggregated AMBCA streamlines showed tangential streamlines (parallel to the cortical surface) in the top and bottom regions of the cortex (**Fig. 5A**, white and yellow arrows). This arraignment was reproduced by the primary orientations of DL-TODs, but not by the primary orientation of CSD-FODs, which showed a uniform radial organization throughout the cortex (**Fig. 5**).

**Fig. 5:**
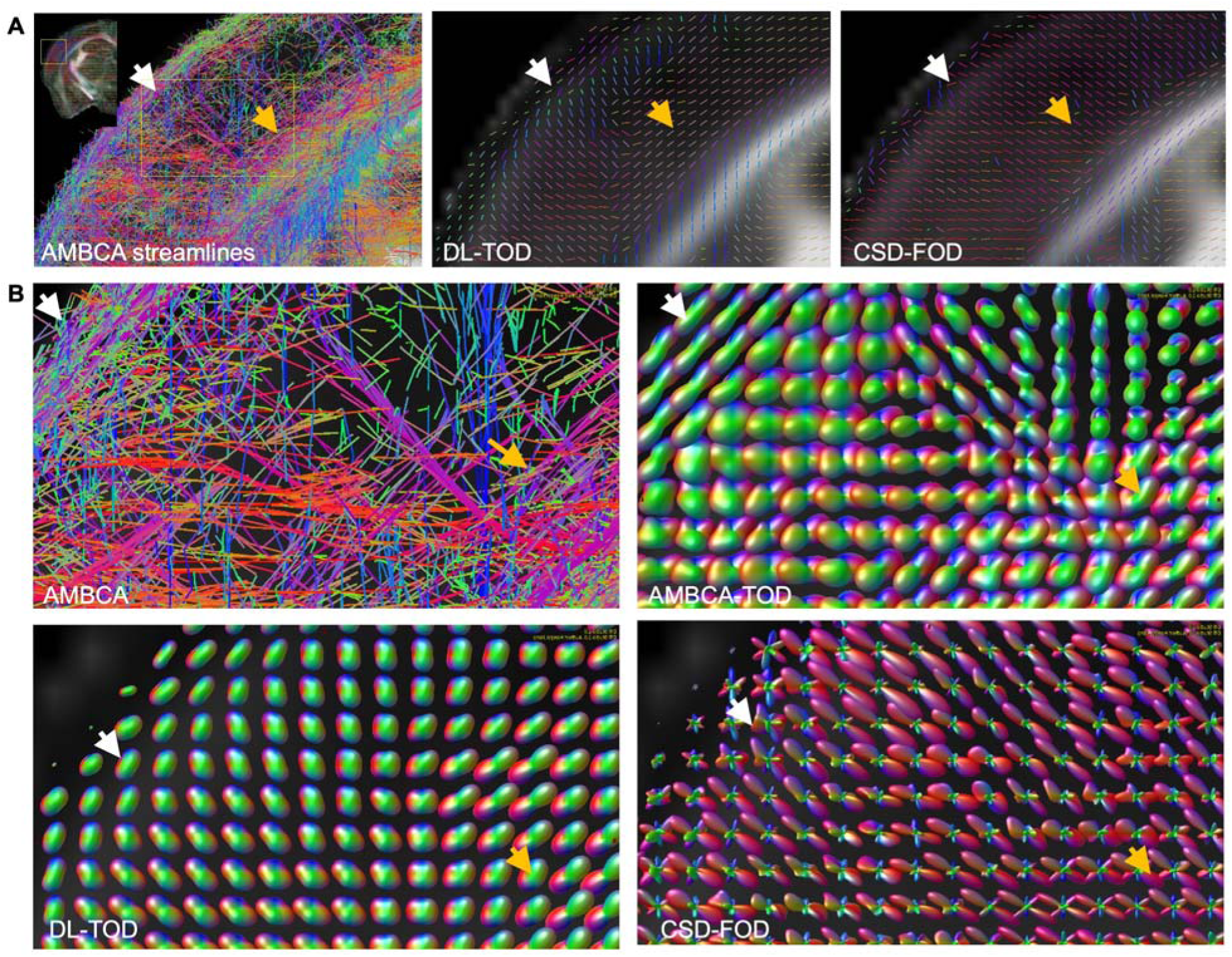
DL-TOD maps recapitulated axonal networks in the cortex. **A:** The aggregated AMBCA streamlines in the sensory cortex (its location is shown in the insert) were compared with the primary orientations of DL-TODs and CSD-FODs. The white and orange arrows indicate the superficial and deep regions of the cortex, where the primary orientation of DL-TODs and CSD-FODs differed. **B:** AMBCA TODs were compared with DL-TODs and CSD-FODs in the same region.

In the hippocampus, the aggregated AMBCA streamlines mostly ran along the left-right (red) or anterior-posterior (green) orientations. The primary orientations of DL-TODs mostly agreed with the AMBCA streamlines (red and green), but the primary orientations of CSD-FODs were along the dorsal-ventral orientation (blue) (**Fig. 6A**). The orientations of the aggregated AMBCA streamlines in the hippocampus were confirmed by the MouseLight dataset. With a few exceptions, axons in the CA1 region of the mouse hippocampus in the MouseLight dataset, either from neurons in the CA1 region or from other parts of the hippocampus, have left-right and anterior-posterior orientations (**Fig. 6C, E-H**), matching the AMBCA streamline results (**Fig. 6A**). In comparison, dense dendritic networks of the CA1 neurons (**Fig. 6D**) from the MouseLight dataset were organized along dorsal-ventral orientation, along the primary directions of CSD-FODs.

**Fig. 6:**
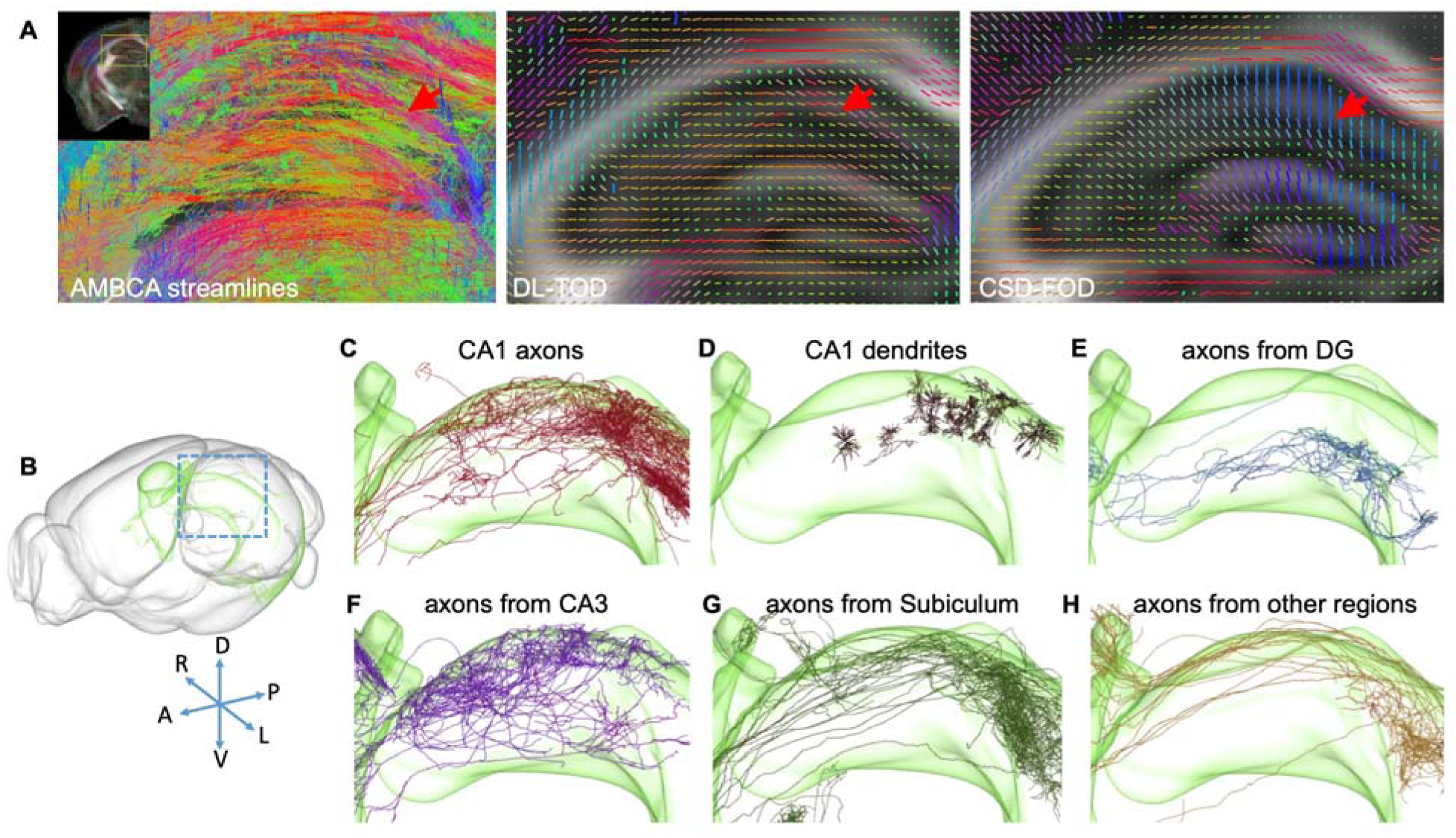
DL-TOD maps recapitulated axonal networks in the hippocampus. **A:** Aggregated AMBCA streamlines in the dorsal hippocampus (its location is shown in the insert) were compared with the primary orientations of DL-TODs and CSD-FODs. The red arrow indicates a group of streamlines running in the horizontal orientation (red), which were found in the primary orientation map of DL-TODs but not in the CSD-FODs. Axons and dendrites in the hippocampus CA1 region from the MouseLight project (Winnubst et al., 2019). **B:** Surface rendering of the hippocampus (green) in the mouse brain. **C-D**: axons and dendrites from CA1 neurons. **E-H:** Axons in the CA1 region but originated from other parts of the hippocampus. DG is the abbreviation of dentate gyrus.

### 3.4 DL-TOD based tractography in the mouse brain

We then examined whether DL-TOD was able to improve tractography in the mouse brain. As explained in section 3.1, the aggregated AMBCA streamlines potentially captured all axonal pathways passing through a region, not just the anterograde pathways originated from it (as in SE AMBCA streamlines). It is therefore appropriate to compare tractography results starting from a seed region with aggregated AMBCA streamlines that passed the same region.

Using a set of ten injection regions in the AMBCA, located throughout the mouse forebrain, streamlines that passed through these regions were selected from the aggregated AMBCA streamlines as well as whole brain tractography results based on CSD-FODs and DL-TODs. Visually, tractography results based on DL-TODs agreed with the selected AMBCA streamlines better than tractography results based on CSD-FODs (**Fig. 7A-C**). We then used the dice index to measure spatial agreements between tractography results and the selected aggregated AMBCA streamlines from the ten regions (**Fig. 7D**). The DL-TOD results had significantly higher DICE scores than the CSD-FOD results in 7 out of the 10 cases and comparable DICE scores for the rest. Further analysis showed that DL-TOD based tractography improved specificity by reducing over-reach, while mostly maintaining a comparable level of sensitivity or overlap compared to FOD-based tractography. In **Fig. 7E**, the DL-TOD-based tractography results, represented by maroon symbols, were mostly to the left of the CSD-FOD-based tractography results, represented by blue symbols. Among the 10 regions, one region in the cortex (ID# 309372716) showed a large drop in sensitivity in DL-TOD-based tractography compared to CSD-FOD based tractography.

**Fig. 7:**
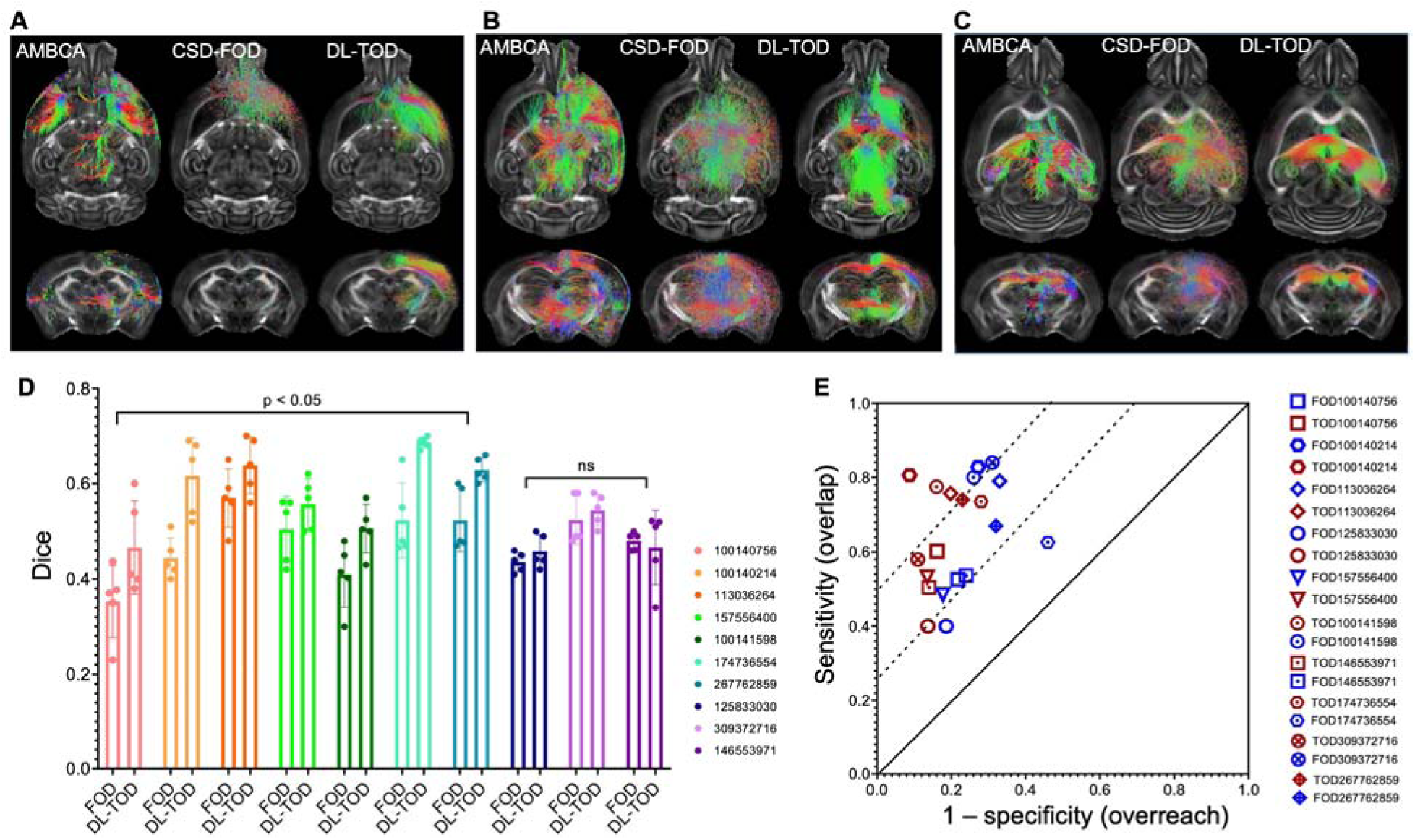
DL-TODs improved dMRI tractography in the mouse brain. **A-C:** Comparisons of streamlines selected from the AMBCA streamline dataset and CSD-FOD/DL-TOD based tractography using injection regions in the frontal cortex (A), thalamus (B), and hippocampus (C). **D:** Spatial overlaps between streamlines selected from the AMBCA streamline dataset and CSD-FOD/DL-TOD based tractography using 10 injection sites in AMBCA measured using DICE. **E:** For the results in E, DL-TOD based tractography results tended to have higher specificity (smaller over-reach) than CSD-FOD based tractography results while maintaining similar sensitivity (overlap).

### 3.5 Potential effects of streamline distribution in AMBCA on DL-TOD

Based on the dMRI signals in the corpus callosum, we simulated dMRI signals from orthogonal crossing fibers, as shown by the estimated CSD-FODs (**Fig. 8A**). In the six-by-six matrix, CSD-FODs along the upper or left edge only had one pair of lobes, representing voxels with a single group of fibers, and CSD-FODs in other places showed two lobes with equal amplitudes, representing the crossing of two groups of fibers. The estimated DL-TODs from the simulated dMRI signals showed less sharply defined lobes than CSD-FODs (**Fig 8A**). When fibers along the x axis (left-right in the mouse brain) were mixed with fibers along the z axis (rostral-caudal in the mouse brain), the estimated DL-TODs showed more preference for the x axis than the z axis, as the DL-TODs in the diagonal elements have their primarily orientations along the x axis. When fibers along the x axis were mixed with fibers along the y axis (dorsal-ventral in the mouse brain), the estimated DL-TODs showed more preference for the y axis than the x axis.

**Fig. 8:**
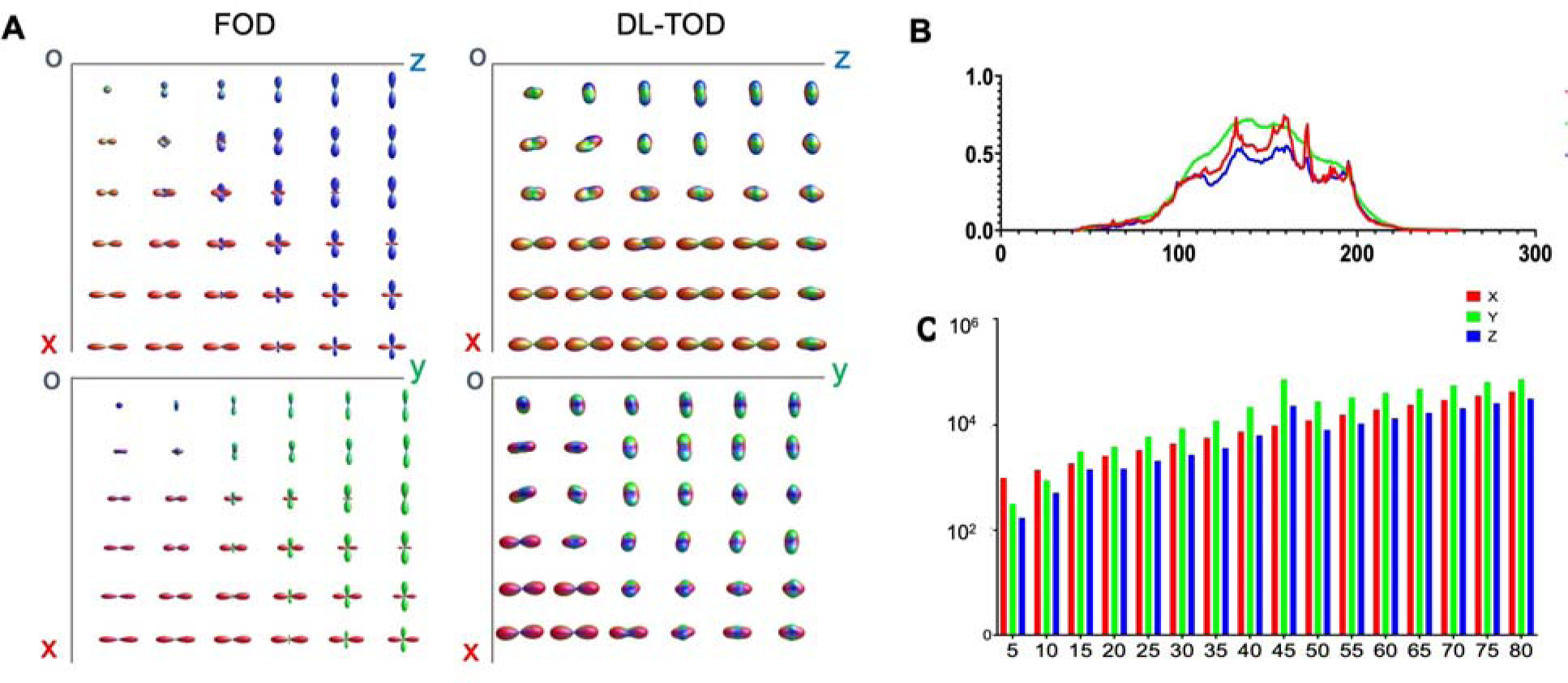
Comparing CSD-FODs and DL-TODs in a digital phantom and potential bias in the AMBCA streamline database. A: Estimated FODs and TODs from a digital phantom containing both single and orthogonal crossing fiber configurations (top: X and Y; bottom: X and Z). **B:** The normalized profiles of AMBCA streamline segments along the X, Y, and Z axes in each axial slice. **C:** The distributions of angles between each AMBCA streamline segments and the three axes.

The preference toward certain orientations in the DL-TODs might reflect the distribution of orientation in the AMBCA streamline dataset. When we projected the orientation of each streamline segment onto the x, y, and z axes, the projections along the y axis out-numbered the projections along the other two axes (**Fig. 8B**), which might have caused the tilt toward the y axis, followed by the x axis, of DL-TODs. Furthermore, examinations of the angles between each streamline segments and the x/y/z axis showed that, among the streamline segments that closely aligned with the x, y, and z axes (angle <10°), there are more streamlines segments closely aligned with the x axis than with the other two axes (**Fig. 8C**). This might explain why the DL-TODs in the simulation were sharper for fiber groups along the x axis than others.

## 4. Discussion

### 4.1 The advantages and limitations of the aggregated AMBCA streamlines

Previous studies on validation and optimization of dMRI based FOD estimation mostly used local microscopy measurements as ground truth (Mollink et al., 2017; Schilling et al., 2016; Schilling et al., 2018). Obtaining comprehensive data on axonal pathways at the mesoscopic or microscopic levels over the entire brain is still technically challenging and time consuming. For example, the connectome of the adult drosophila brain was completed only recently (Scheffer et al., 2020). The aggregated AMBCA streamlines, which combine streamlines from multiple experiments and are therefore more comprehensive than individual tract tracing experiments, may be a suitable candidate for validation and optimization of FOD estimation and dMRI tractography. Although it would be ideal to have dMRI and tract tracing data from the same animals as suggested by previous studies (Grisot et al., 2021), it is impractical to re-acquire a comprehensive collections of tracer results using methods based on viral or chemical tracers. This study was therefore based on two assumptions: 1) the aggregated AMBCA streamlines, with data from more than 2,700 subjects, captured common forebrain axonal pathways in the C57BL/6 mouse forebrain; and 2) the inter-subject differences among the inbred C57BL mouse brains were small so that the aggregated AMBCA streamlines can be used as the ground truth to compare with dMRI signals acquired from a separate cohort of C57BL/6 mice at the same age as the mouse brains in AMBCA.

Although it is difficult to confirm the first assumption, evidence from comparing the aggregated AMBCA streamlines with cell tracing results from the MouseLight projects (**Fig. 3B-C**) suggests that at least the axonal pathways in the cortex and hippocampus were well covered. The aggregated AMBCA streamlines, however, remained an incomplete representation of the axonal networks in the mouse brain in several other aspects due to several limitations, including non-uniform distribution of injection sites, varying dosages and labelling efficiency, imaging resolution, and the fast-marching algorithm. For example, as the cerebellum received fewer injections than the cortex, the density of streamlines in the cerebellum was noticeably lower than in the forebrain as shown in Fig. 2D. Due to this reason, although the orientations of TODs should mostly reflect those of actual axonal pathways, the amplitudes of the AMBCA TODs were not used in this study. This limitation may be alleviated with more tract and cell tracing resources in the future (e.g. (Winnubst et al., 2019; Zingg et al., 2014)). It is also not clear whether the technique developed using the aggregated AMBCA streamlines can be translated to data from the human brain, due to the vast differences in imaging resolution as well as differences in microstructural organization. The recent reports of tract tracing in non-human primate brain (Charvet et al., 2022; Howard et al., 2019; Xu et al., 2021) offer hope that such data, which have higher translational value, may become more readily available in the future. For the second assumption to hold, it is necessary to image a large number of subjects, more than the number of subjects in this study so that impacts of individual variations in microstructural organization including axonal pathways will be reduced.

### 4.2. Deep learning can enhance the estimation of FOD

The deep learning framework has several advantages that complement existing modeling approaches, as it is data-driven and not limited by underlying assumptions associated with models. Here, we trained out network using small 3x3x3 patches instead of entire images because we assumed the relationship between dMRI signals and underlying axonal pathways to be strictly local (i.e. MRI signals are the ensemble average of spins within each voxel and do not depend on neighboring voxels) and would like to limit the amount of spatial information available to the network, in a similar fashion as our previous study connecting MRI signals and histology (Liang et al., 2022). As a result, a limited number of typical dMRI data, instead of thousands as required in typical deep learning studies, can provides sufficient instances to train the deep learning network to resolve the orientation of certain axonal pathways in gray matter. The network can be extended to include dMRI data acquired using optimized parameters or other MRI contrasts to further enhance its capability. For example, the recent developments on the multi-dimensional diffusion encoding (Topgaard, 2017) may provide critical information on tissue microstructural organization that can be utilized by the network.

Our results demonstrated that deep learning networks, trained with ground truth tract tracing data, were able to improve the estimation of FODs (**Figs. 5-6**) and tractography (**Fig. 7**) in the mouse brain. The improvements were most obvious in gray matter regions (**Fig. 4B-C**) since conventional CSD-FODs are already fairly accurate in depicting the orientation of white matter bundles (Lin et al., 2001; Mollink et al., 2017; Schilling et al., 2018). The complex microstructural organization in gray matter, with the presence of dendritic networks and other cellular compartments, however, challenges the power of dMRI in resolving axonal pathways in gray matter regions, as shown previously in the mouse hippocampus (Wu and Zhang, 2016) and other regions (Moldrich et al., 2010; Ren et al., 2007; Wu et al., 2014). The deep learning network developed here, as shown in **Fig. 6**, may be used to reconstruct structural connectivity in the mouse hippocampus or other regions.

One challenge introduced by AMBCA TODs was that, while AMBCA TODs in large white matter tracts had sharp lobes that aligned with CSD-FODs, AMBCA TODs in gray matter regions (e.g. the cortex) tended to have less well-defined FOD lobes (**Figs. 4 and 5**). This potentially reflected the large dispersion of streamlines in these regions, which is different from the assumption behind the estimation of CSD-FODs using a single response function based on dMRI signals from white matter structures. We did not compare our results with recent work on developing more accurate estimation of FODs in gray matter region (De Luca et al., 2020; Jeurissen et al., 2014), which should be considered for future studies.

The deep leaning method used in our work still has room for improvement. For example, the MSE loss function used in training the network may not be optimal as it assigned equal weighting to all 28 TOD spherical harmonic coefficients. As some TOD coefficients can have much higher amplitudes than others, the loss function used here may have introduced a bias for high amplitude coefficients. One solution is to use multiple loss functions (Ghodrati et al., 2019; Kendall et al., 2018; Wang et al., 2022), which should be explored in future studies. Another limitation of the deep learning approach here is that biases in the training data will influence the network predictions. The mouse brain has a higher relative gray matter volume than the human brain. In addition, the distribution of streamlines’ orientation, as shown in **Fig. 8B**, was not uniform, which may reflect actual axonal organization in the brain (i.e. more axons running horizontally than axons running vertically) or may be a result of the atlas space used here. In the future, training datasets containing with equalized tissue types and multiple orientations may be able to reduce the bias.

Our results also did not provide detailed knowledge of the inner working of the network. For example, it is intriguing that the network was able to assign distinct orientations to the superficial, middle, and deep cortical regions based on dMRI signals (**Fig. 5**). The normalized dMRI signals in these three regions and corpus callosum are displayed in **Fig. 9**. The dMRI signals in the corpus callosum had less attenuations than signals from the three cortical regions, and stronger signal attenuations were observed when the diffusion encoding gradients were close to the left-right direction (polar angle close to 0 and ±π in the plot) than otherwise. The dMRI signals from the three cortical regions all showed higher degrees of attenuations than signals from the corpus callosum but displayed different patterns. All three regions showed strong signal attenuations in region 2 in **Fig. 9**, but the superficial and deep cortical regions had less attenuation than the middle cortical region in region 2, and much less attenuation in regions 1 and 3, potentially due to different fiber orientations. The deep learning network might have recognized the differences in dMRI signals among the three regions and used the information to predict the DL-TOD, whereas the conventional CSD methods using a single response function derived from white matter regions did not resolve the fiber orientations accurately in this case.

**Fig. 9:**
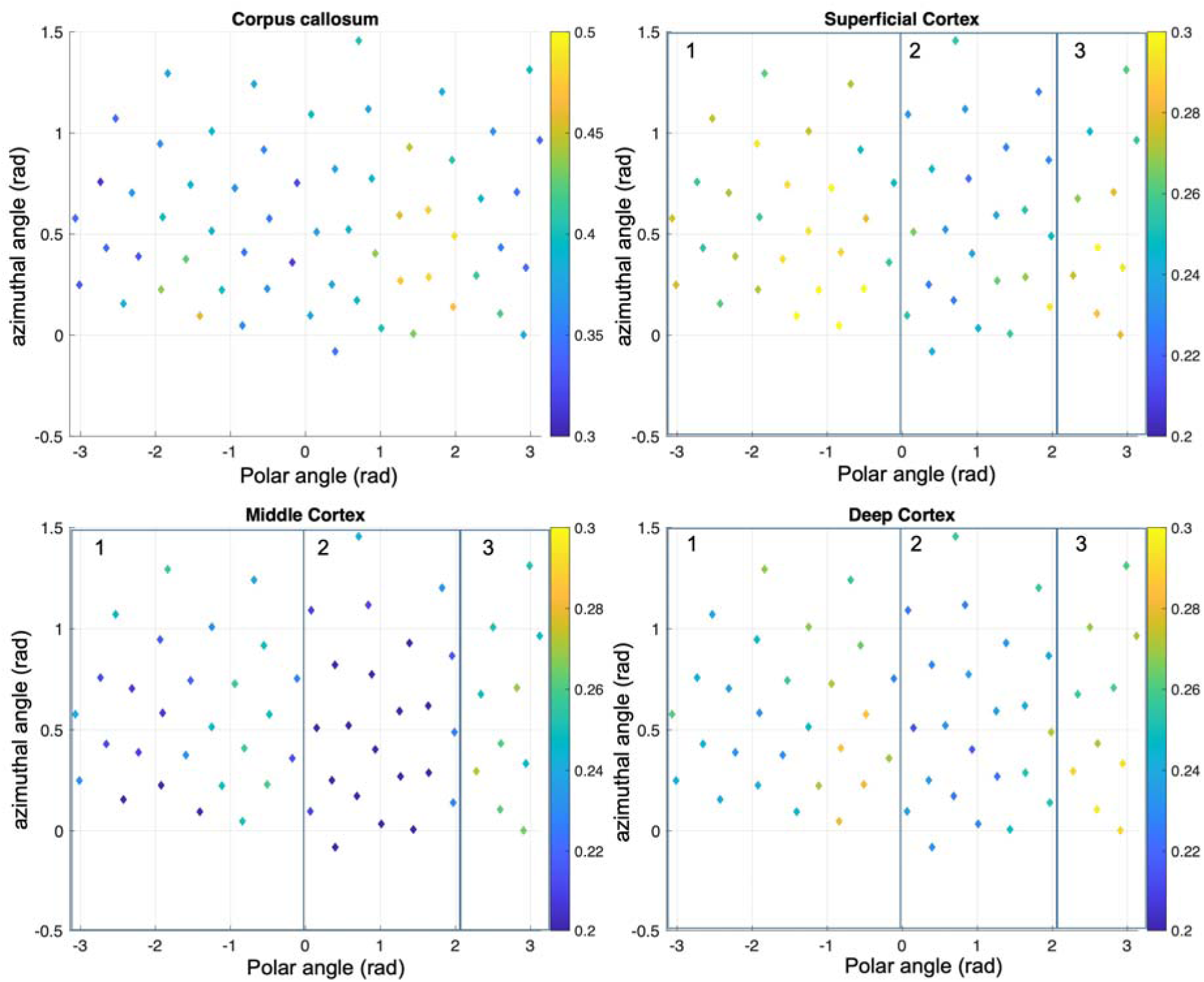
Normalized dMRI signals from representative voxels in the corpus callosum, superficial, middle, and deep cortical regions. The locations of the signals in the plot indicate the polar and azimuthal angles of the corresponding diffusion encoding gradient, and the color indicates the normalized signal strength (with respect to non-diffusion-weighted signals). The plots of cortical dMRI signals are overlaid with three rectangular boxes to guide visual comparisons.

In summary, we demonstrated that a deep learning network, trained using the aggregated AMBCA streamline, can improve the estimation of FODs as well as the specificity of tractography from mouse brain dMRI data. The technique developed here can be used to characterize structural connectivity in the mouse brain.

## Acknowledgement

This work was supported by NIH grants R01NS102904 and R01HD074593. The majority of this work was performed at the NYU Langone Health Preclinical Imaging Laboratory, a shared resource partially supported by the NIH/SIG 1S10OD018337-01, the Laura and Isaac Perlmutter Cancer Center Support Grant, NIH/NCI 5P30CA016087, and the NIBIB Biomedical Technology Re-source Center Grant NIH P41 EB017183 as well as by the NYU CTSA grant UL1 TR000038 from the National Center for Advancing Translational Sciences, National Institutes of Health.

## Data and materials availability

The analyses in this study were carried out on publicly available datasets. The mouse brain streamline data were obtained from the Allen mouse connectivity project (http://connectivity.brain-map.org). Code and data for training are available at https://github.com/liangzifei/MR-TOD-net. The aggregated AMBCA streamlines can be downloaded from https://osf.io/m98wn.

